# Successful reproduction of a large EEG study across software packages

**DOI:** 10.1101/2022.08.03.502683

**Authors:** Aya Kabbara, Nina Forde, Camille Maumet, Mahmoud Hassan

## Abstract

As an active field of research and with the development of state-of-the-art algorithms to analyze EEG datasets, the parametrization of Electroencephalography (EEG) analysis workflows has become increasingly flexible and complex, with a great variety of methodological options and tools to be selected at each step. This high analytical flexibility can be problematic as it can yield to variability in research outcomes. Therefore, growing attention has been recently paid to understand the potential impact of different methodological decisions on the reproducibility of results.

In this paper, we aim to examine how sensitive the results of EEG analyses are to variations in preprocessing with different software tools. We reanalyzed the shared EEG data (N=500) from (Williams et al. 2021) using three of the most commonly used EEG software tools: EEGLAB, Brainstorm and FieldTrip. After reproducing the same original preprocessing workflow in each software, the resulting evoked-related potentials (ERPs) were qualitatively and quantitatively compared in order to examine the degree of consistency/discrepancy between softwares. Our findings show a good degree of convergence in terms of the general profile of ERP waveforms, peak latencies and effect size estimates related to specific signal features. However, considerable variability was also observed in the magnitude of the absolute voltage observed with each software package as reflected by the similarity values and observed statistical differences at particular channels and time instants. In conclusion, we believe that this study provides valuable clues to better understand the impact of the software tool on the analysis of EEG results.

## 1. Introduction

Electroencephalography (EEG) is a well-established technique for measuring the electrical fluctuations generated by the brain at high temporal resolution. Due to its non-invasiveness, low cost and ease-of-use EEG has been gaining increasing interest in uncovering the functional brain activity underlying various brain conditions including disorders, emotions, information processing and resting state (Lopes da Silva 2013).

Typically, the EEG electrodes capture a mixture of neural activity and non-neural-related artifacts which can be physiological (e.g. eye movements or muscle contractions) or external to the human body (e.g. power line or interference with other electrical devices) (Urigüen and Garcia-Zapirain 2015). Thus, to study the EEG signal, it is of great importance to first carefully reduce the influence of contaminating artifacts while preserving the neural activity. This is the aim of the preprocessing stage which is carried out so as to derive clean EEG signals suitable for further statistical analysis. Preprocessing typically includes multiple steps, such as line noise removal, re-referencing, artifact rejection, filtering, epoch selection, bad channels detection and interpolation. Although there is a general agreement in the scientific community on the main steps that should be considered in the preprocessing pipeline, each step can be approached through many algorithmic strategies with different sets of assumptions, and parameter choices (Šoškić et al. 2022; Boudewyn et al. 2018; Croft et al. 2005; Šoškić et al. 2021). Thus, the preprocessed signals are the result of multiple individual and user-dependent decisions, made over a potentially long and ordered pipeline. More specifically, the chain of decisions is not only limited to adjusting the features incorporated in each preprocessing step, but may even start before the preprocessing is performed - i.e, when selecting the adequate software tool.

In this context, many efforts have focused on proposing guidelines for researchers to choose between the existing cleaning methods depending on the application and user’s requirements. Among these efforts, (Islam, Rastegarnia, and Yang 2016; Ranjan, Chandra Sahana, and Kumar Bhandari 2021) present extensive reviews of the existing state-of-the-art artifact cleaning methods by showing the pros, cons and suitability in particular applications.

Recently, growing attention has been paid to evaluate the variability and comparability of results obtained with different preprocessing methods and parameter choices (Barban et al. 2021; Robbins et al. 2020; Clayson et al. 2021). The main objective of these studies was to test how much the variability in cleaning methods can impact the conclusions of a study. For instance, the effect of three artifact removal algorithms (ICA-LARA, ICA-MARA and Artifact Subspace reconstruction (ASR)) on EEG characteristics and event-related measures was analyzed and compared across 17 EEG studies (Robbins et al. 2020). Results highlight the existence of significant differences between results particularly after eye blinks artifacts have been removed. Others were interested in testing the ability of different blind source separation methods to remove synthetic/modeled noise sources corrupting real EEG signals (Barban et al. 2021). The main results show that there is no method that can be considered as an all-purpose algorithm, and the choice of the adopted method should be driven by the specific needs of users (such as the computational capacities, or the temporal constraints). Trying to optimize the preprocessing pipeline for the event-related potentials (ERPs), (Clayson et al. 2021; Šoškić et al. 2022) examined the impact of many possible methodological choices on the data quality and the experimental effects through data multiverse analysis. Both studies highlighted the substantial impact of several parameters such as the filter cut-off, artifact detection method, baseline adjustment, reference, scoring electrodes and others on the study outcomes.

While the above studies provide important insights on the effect of either the preprocessing stages, the preprocessing algorithms or the parameter choices, the preprocessing of the signals were carried out using a single software tool. Yet, there are many tools available to study the EEG signal including: EEGLAB (Delorme and Makeig 2004), Brainstorm (Tadel et al. 2011), MNE (Gramfort et al. 2014), FieldTrip (Oostenveld et al. 2011) and Automagic (Pedroni, Bahreini, and Langer 2019). Each toolbox has its own way to organize and format the data, to implement functions and to define their arguments, parameters, optimal and default values. Another important difference between tools resides in the availability of the desired preprocessing steps as well as the parameters that can be accessed for each step.

Here we investigate how sensitive the results of EEG analyses are to variations in software packages when using the same dataset and aligned preprocessing methods. To this aim, we reanalyzed data (N=500) from a recent study by Williams and colleagues (Williams et al. 2021) and reproduced the study using three software packages to quantify the observed differences in the final results. Our objective was first to reproduce the main figures of (Williams et al. 2021) by replicating the original preprocessing pipeline used within each software tool. We compare three of the most commonly used MATLAB toolboxes: Brainstorm, EEGLAB and FieldTrip in order to achieve two main objectives: 1) Study whether the main findings of the original paper – including ERP curve as well as effect size estimates related to selected ERP features – could be reproduced within each software packages. 2) Quantify variations observed across software packages.

## 2. Materials and methods

### 2.1. Material

#### 2.1.1. Dataset

We used the dataset previously analyzed in (Williams et al. 2021), and publicly available at www.osf.io/65x4v/. In brief, this dataset comprises data from 500 undergraduate healthy students (341 female, mean age=21.71 years old, 440 right handed) recruited by the University of Victorial. These participants were selected amongst a total of 637 subjects as they had provided signals with a high data quality. The study was approved by the University of Victoria’s Human Research Ethics Board and all participants provided written informed consent before any data acquisition.

We chose to reproduce this study by Williams and colleagues for two main reasons. First the availability of the raw data and of the preprocessing and analysis scripts made it possible for us to recompute the original results to serve as a reference for our subsequent analyses Second, the large number of participants (N=500) made this study less sensitive to a lack of reproducibility that would be due to small sample size (Ioannidis 2005; Button et al. 2013).

#### 2.1.2. Experimental protocol

Participants completed a simple gambling task following a two-armed bandit task. This task was chosen by Williams and colleagues in (Williams et al. 2021), as it is the most commonly used paradigm to evoke the reward positivity ERP which was the subject of investigation of the reference paper (Proudfit 2015). The pipeline of our study is summarized in Figure 1.

**Figure 1.**
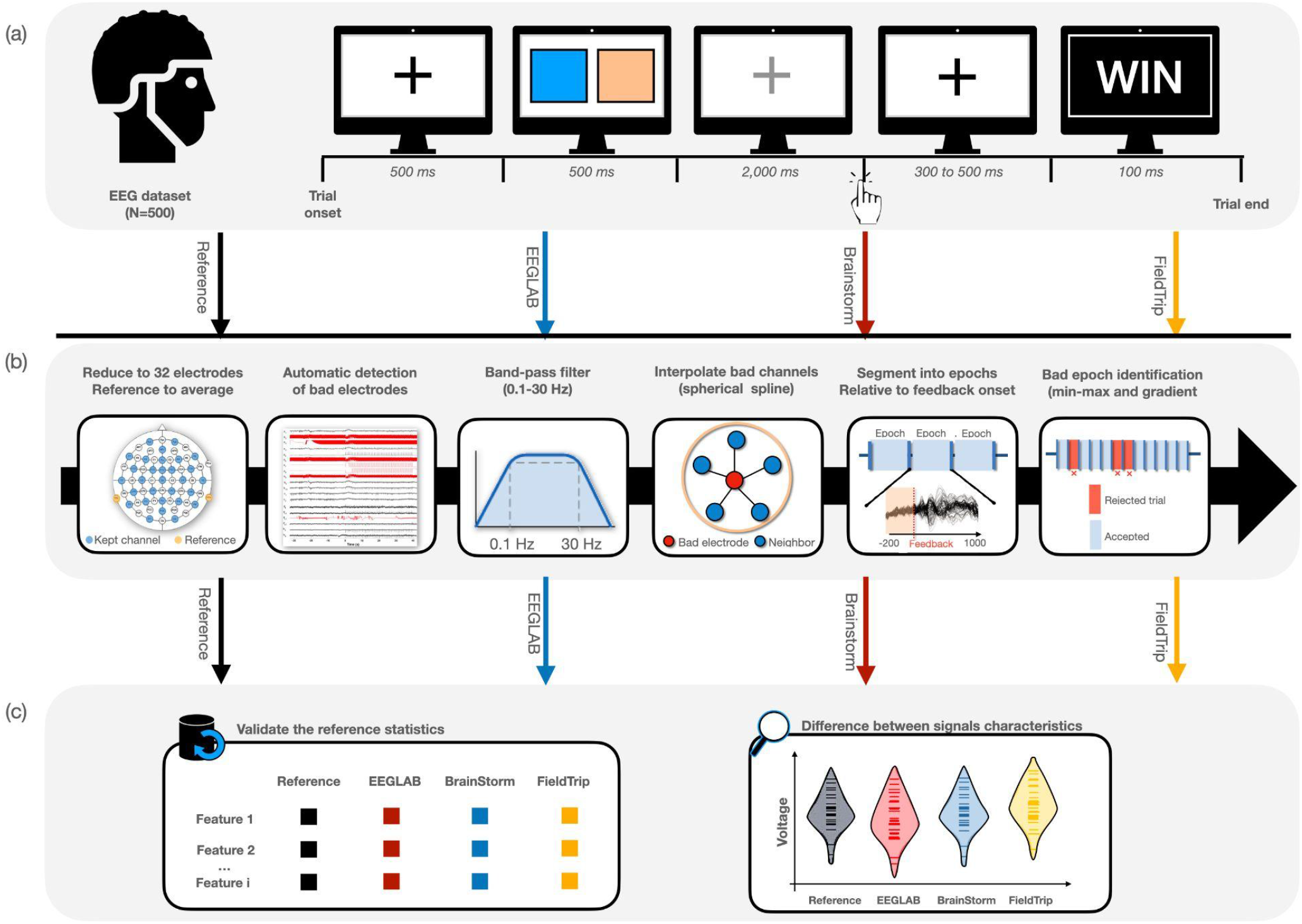
Overview of the study. (a) We used shared EEG data from (Williams et al. 2021) with 500 participants performing a simple gambling task of six blocks composed of 20 trials. (b) This dataset was then preprocessed using the different software tools: Reference (using the code published with the original paper), EEGLAB, Brainstorm and FieldTrip. The preprocessing steps to be performed in each tool included: reduction to 32 electrodes, re-referencing, automatic detection of bad electrodes, band-pass filtering, interpolation of bad channels, segmentation into time-locked epochs and removal of artifactual trials. (c) The preprocessed signals derived from the four preprocessing codes were used to reproduce the reference statistics and hypotheses. A quantitative comparison between the resulting signals was also conducted in terms of signal features. Image credits: EEG cap CC-BY Wikimedia Commons by CIV The Noun Project.

The acquisition session consisted of six blocks of 20 trials (see Fig. 1.a). Each trial was initiated by a black fixation cross displayed for 500 ms, followed by a 500 ms display of two colored squares. Then, the fixation cross turned gray to prompt the participant to select one of the two colored squares (left or right) within a 2,000 ms time limit. After that, a black fixation cross was presented for 300 to 500 ms, and a simple feedback (“WIN” for gain, “LOSE” for loss) was shown for 1,000 ms. The final objective of this task for the participant was to win as often as possible. This was possible for the participant by determining (while computing the task) which square would bring the most successful rate (60% for one square vs. 10% for the other one). The same pair of colors was used for all the trials of the same block, and the squares locations were randomized for each trial.

#### 2.1.3. Data collection

EEG data were acquired from either 64 or 32 electrode (Ag/AgCl) EEG systems (ActiCAP, Brain Products, GmbH, Munich, Germany) using Brain Vision Recorder. Data were originally sampled at 500 Hz and low-pass filtered below 245 Hz. During the recording, all electrodes impedance were kept under 20 kΩ on average.

### 2.2. Methods

#### 2.2.1. Original preprocessing pipeline

The preprocessing pipeline adapted by the reference paper was performed in MATLAB using scripts available at www.osf.io/65x4v/ (the main file is named *‘RewardProcessing_Preprocessing*.*m’*), where some functions have EEGLAB dependencies. The pipeline consisted of multiple steps (Figure 1.B) as follows:

1. **Reduce the number of electrodes to 32** electrodes (for all data that were collected with a 64 electrode EEG system).
2. **Detect artifactual channels** Practically, the detection of artifactual channels can be approached in different ways. Among these strategies, Williams and colleagues used the trial rejection rate. The data first were re-referenced to an average mastoid reference (using TP9 and TP10 electrodes) and band pass-filtered between 0.1 to 30 Hz (Butterworth, order 4). A notch filter at 60 Hz was also applied. Afterwards, authors have corrected eye blinks after manually identifying the corresponding independent components (ICs) reflective of blinks. Time-locked epochs around the feedback stimulus onset (from -500 to 1500 ms) were then extracted, and baseline corrected using a −200 to 0 ms window. An artifactual trial (i.e epoch) was identified with 10 μV/ ms gradient and 100 μV maximum–minimum criteria. Ultimately, an electrode was considered noisy or artifactual if it exceeded a trial rejection rate of 40%.
3. **Re-reference data to an average mastoid reference** (using TP9 and TP10 electrodes)
4. **Apply a band pass-filter** between 0.1 to 30 Hz (Butterworth, order 4) and a **notch filter** at 60 Hz.
5. **Interpolate the detected artifactual channels** using the spherical spline method.
6. **Detect and remove the eye blinks artifacts** using independent component analysis (ICA) after manually selecting the blinks components via topographic maps and component loadings.
7. **Extract the time-locked events** using a segment window of −500 to 1,300 ms relative to the feedback stimulus.
8. **Remove baseline values** using a window of -200 to 0 ms.
9. **Reject trials** that exceed a gradient of 10 μV/ ms and a maximum-minimum voltage of 100 μV.
10. **Compute the ERPs of ‘Win’ and ‘Loss’ conditions** by averaging the corresponding epochs and ERPs were trimmed to −200 to 1,000 ms. Authors were also interested in analyzing the grand averaged ERP, denoted the reward positivity, obtained as the result of subtraction between the gain condition and the loss condition.

In the original preprocessing pipeline proposed by the authors, a manual procedure – i.e., a human-based and visually guided procedure – was used to detect the components corresponding to the eye blinking noise. However, this step is not only time-consuming to be carried out in each software tool for 500 subjects, but more importantly also introduces inter-rater variability as it is open to the level of expertise and variability across different raters performing the manual detection. In order to focus on inter-software variability only, this step was removed from the preprocessing pipeline.

#### 2.2.2. Comparison across toolboxes

We selected three of the most widely used toolboxes available to preprocess EEGs and reproduced the reference preprocessing pipeline in each.

All code to reproduce the preprocessing pipelines is available at: https://github.com/Inria-Empenn/EEG_preprocessing (released on Zenodo, doi: 10.5281/zenodo.6918329) and more details are provided below on the algorithms and parameters chosen in each software package.

##### 2.2.2.1. EEGLAB

The EEGLAB preprocessing script was assembled and run for all the 500 subjects as follows:

1. Load the data using *pop_loadbv*.*m*
2. Reduce data into 32 channels using *pop_select*.*m*
3. Automatically detect the noisy channels: Given that the procedure proposed by the reference paper was not available in EEGLAB, we relied on EEGLAB automatic artifactual channel detection. In practice, EEGLAB incorporates a plugin named *‘clean_rawdata’* that detects flatline and noisy channels based on three criteria: i) the channel has no signal variation for a duration of longer than a specific time window length (default 5s), ii) the channel presents a correlation to its robust estimate (based on other channels) lower than a given threshold (default 0.85), iii) the channel presents an excessive line noise by exceeding a threshold value in standard deviations above the channel population mean (default to 4 standard deviations).
4. Interpolate the detected noisy channels using the spherical spline method *pop_interp*.*m*
5. Re-reference the signals to the average mastoid electrodes using *prop_reref*.*m*.
6. Filter the signals between 0.1 and 30 Hz using *pop_eegfiltnew*.*m*. This function uses a hamming window-based finite impulse response (FIR) filter.
7. Divide the signals into time-locked epochs using the function *pop_epoch*.*m*, and correct to baseline using *pop_rmbase*.*m* function
8. Reject the artifactual trials using the *pop_eegthresh*.*m* function for which the lower and upper amplitude limits can be identified by the user. Here, we set the lower limit to -50μV and the upper limit to 50μV, in a way to follow the same parameters of the trial rejection procedure adopted in the reference paper.
9. Compute the conditional ERPs (for win and loss conditions) as well as the grand averaged ERP

The analysis was conducted using EEGLAB v2021.1 (RRID: SCR_007292).

##### 2.2.2.2. Brainstorm

The list of steps used to perform the preprocessing in Brainstorm are as follows:

1. Re-reference to the average mastoid (using TP9 and TP10) using *Process_eegref*
2. Detect the noisy channels using *Process_detectbad*. It is noteworthy to mention that Brainstorm does not provide an automatic approach to detect the noisy channels. Instead, this detection is typically done manually by visual inspection. To automate the detection of flat channels, *process_detectbad* can detect the bad channels using a peak-to-peak threshold criteria.
3. Interpolate the detected noisy channels using the spherical spline method with *Process_eeg_interpbad*.
4. Apply the notch filter at 60 Hz using *Process_notch*.
5. Apply a band-pass filter between 0.1 and 30 Hz with a linear phase FIR filter using the process *Process_bandpass*.
6. Segment data into time-locked epochs using *Process_import_data_event*.
7. Remove baseline values from -200 ms to 0 using *Process_baseline*.
8. Detect and reject the bad trials using a peak to peak of 100 microVolts using *Process_detectbad*.
9. Compute the conditional ERPs using *Process_average*

The analysis was conducted using brainstorm version 20.09.28 (RRID: SCR_001761).

##### 2.2.2.3. FieldTrip

FieldTrip toolbox is not a software with a user interface, but rather a collection of functions. Thus, a Matlab script, in which a sequence of FieldTrip functions are called, is considered as an analysis protocol in FieldTrip. Each of the functions of the toolbox takes as input the data that was produced by the previous function. To allow a function to implement a specific algorithm, particular parameters can be specified via a configuration structure *cfg*. Here, we used the major functions *ft_preprocessing, ft_artifact_clip, ft_channelrepair, ft_redefinetrial, ft_rejectartifact* and *ft_timelockanalysis*.

More precisely, the FieldTrip preprocessing script was assembled and run for all the 500 subjects as follows:

1. Data were first reduced to 32 channels, re-referenced to the average mastoid, filtered by a butterwoth filter (order=4) using *ft_preprocessing* with *cfg*.*channel, cfg*.*refchannel, cfg*.*bpfreq, cfg*.*bpfilttype, cfg*.*bpfiltord* being adequately defined.
2. Bad channels were detected as those showing signals being completely flat for a given time window. Here, we set the length of the time window to 5s. The function used is *ft_artifact_clip* where *cfg*.*artfctdef*.*clip*.*timethreshold* was defined.
3. The interpolation of the detected bad channels was done using *ft_channelrepair*, with *cfg*.*badchannel* being identified.
4. The segmentation into time-locked epochs to win and loss conditions was performed using *ft_redefinetrial* where *cfg*.*trialdef* is configured.
5. A correction to baseline was performed using *ft_preprocessing* after defining *cfg*.*baselinewindow*.
6. A trial is detected as bad using *ft_artifact_threshold* if it exceeds a min-max voltage of 100 microVolts following the same criteria of the reference paper, then rejected using *ft_rejectartifact*.
7. The conditional ERPs were computed using *ft_timelockanalysis*.

The analysis was conducted using FieldTrip version 20191024 (RRID: SCR_004849).

#### 2.2.3. Reproduction of the ERP analysis

In (Williams et al. 2021), the authors focused on analyzing the neural feedback processing based on multiple measures of reward positivity. Many of these measures rely on the ERP, which attempts to characterize the neural activity by examining the peaks and troughs of the averaged signals time-locked to events of interest (Picton, Lins, and Scherg 1995). More specifically, the authors have first computed the conditional ERPs for each condition (gain and loss) within each participant. Difference ERPs were also extracted by subtracting the conditional ERP related to the loss condition from that related to the gain condition. Then, four quantitative ERP-based features were determined corresponding to FCz electrode,the most commonly used electrode in the context of reward positivity (Sambrook and Goslin 2015): *Peak time of the reward positivity*: computed for each participant by finding the peak amplitude of the difference ERP waveform. *Mean peak*: Average of the voltages ±46 ms surrounding the peak location. *Maximum peak*: Largest amplitude within the 200 to 400 ms time window. *Base-to-peak*: Measure computed by subtracting the minimum voltage of the trough immediately prior to the reward positivity from the maximum peak measure.

The mean, maximum and base-to-peak metrics were computed for both the conditional and the difference ERPs of each participant. In our study, we followed the same ERP exploration and features extraction procedures after obtaining the preprocessed signals from the different softwares.

#### 2.3. Comparison methods

We applied three separate quantitative methods to measure the discrepancy between the results obtained within each software. First, the statistical comparisons among metrics (mean, maximum, base-to-peak and peak-time) obtained by the different software tools were performed using Wilcoxon ranksum test. For each metric of interest, we compared the values obtained by the reference, EEGLAB, BrainStorm and FieldTrip for all participants. These comparisons provide a quantification of the level of (dis)agreement between each pair of software tools about the ERP features of interest. The statistical significance level was set to *p* < 0.01 and Bonferroni correction was used to address the multiple comparisons issue across the number of tests performed (6 comparisons).

Second, we evaluated the variability of results by computing the similarity between the ERPs obtained by the different software tools, when considering all the EEG channels. In fact, in their paper, Williams and colleagues have only considered the FCz electrode as it was shown to be the electrode that extracts the most relevant information related to their topic of interest (i.e the reward positivity). Here, however, we are also interested in studying the effect of the software tool on the preprocessed EEG signals of all the recording channels. Thus, for each participant, we assessed the similarity between two softwares *S1* and *S2* using pearson’s correlation measure as follows:

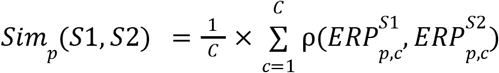

Where *p* is the considered participant, *C* is the number of channels. 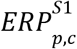 and 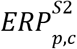 denote the ERP signal obtained at channel c from the software S1 and the software S2, respectively.

In addition, we were interested in precisely describing the ERP differences between softwares in terms of temporal and spatial characteristics. This was done by assessing the statistical difference between ERP distributions at each millisecond and each channel using wilcoxon ranksum test, which yields a 2D map of p-values between each two softwares. To account for the multiple comparisons problem, all p-values were corrected across all timepoints and software studied using Bonferroni resulting in a thresholding p-value of ^−7^

### 2.4. Code availability

Codes supporting the results of this study are available at https://github.com/Inria-Empenn/EEG_preprocessing (released on Zenodo, doi: 10.5281/zenodo.6918329). All the preprocessing codes were written in Matlab (Matlab 2018). The visualizations of ERP waveforms (Figure 2) and the quantitative features (Figure 3) were done in R (R Core Team 2020). Seaborn (Waskom 2021) was used to illustrate the comparisons between the software distribution of the quantitative measures (Figure 4), and the similarity matrix between softwares (Figure 5). Other visualizations and statistical assessments were conducted using Matlab.

**Figure 2.**
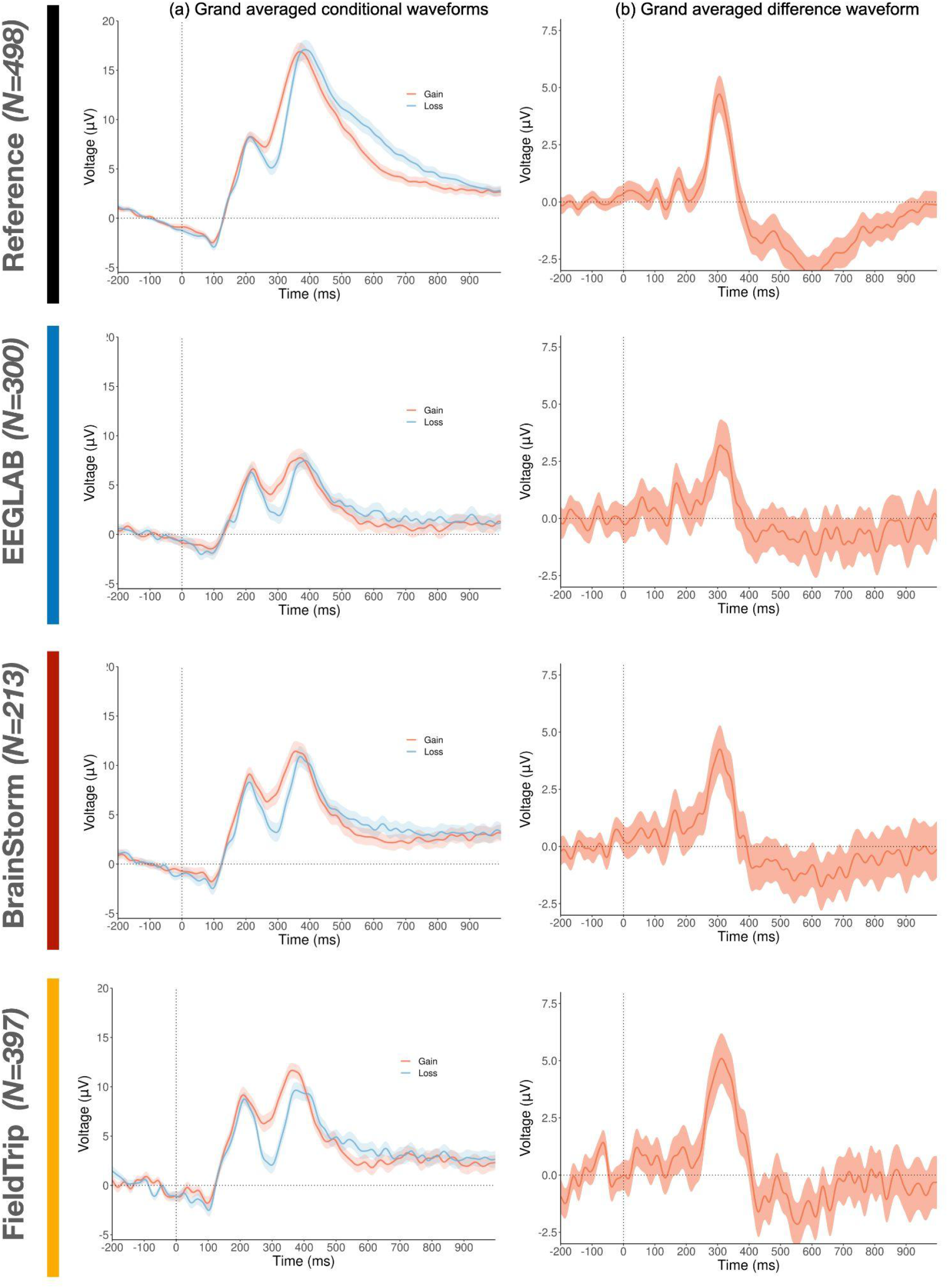
ERP waveforms at electrode FCz illustrating the reward positivity after preprocessing by: the reference code, EEGLAB, Brainstorm and FieldTrip. (a) grand averaged conditional waveforms (ERP averaged across all subjects) with 95% confidence intervals, (b) grand averaged difference waveform with 95% confidence intervals. These subfigures are reproduced from Figure 3 illustrated in (Williams et al. 2021).

**Figure 3.**
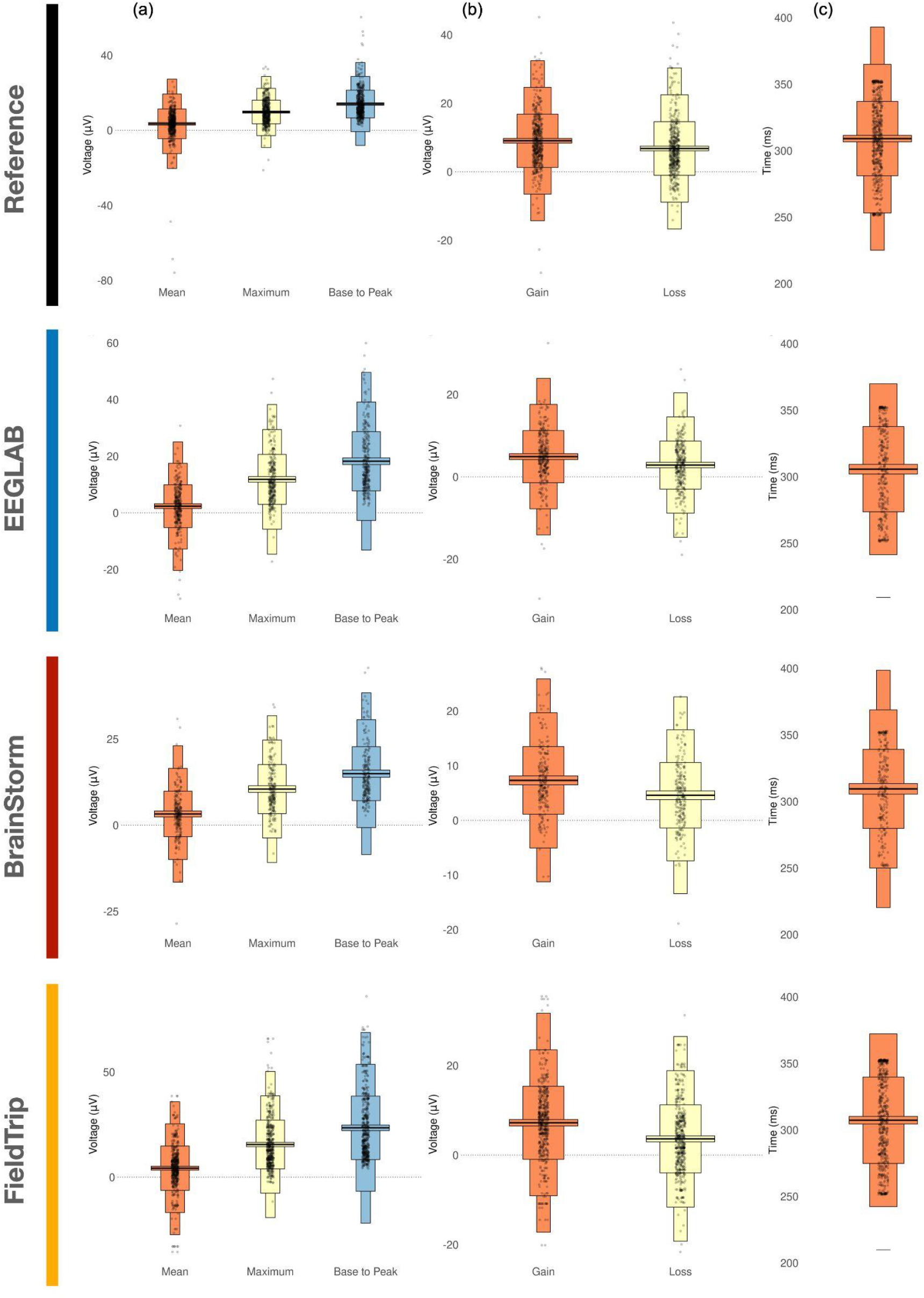
The metrics distribution across all participants for the different preprocessing softwares. (a) The features calculated on the difference ERP, (b) conditional amplitudes for the mean peak measure, and (c) peak latency of the reward positivity (difference ERP). Each black dot represents a participant’s data and the middle black lines represent the mean across participants. These subfigures are a reproduction of Figure 3 illustrated in (Williams et al. 2021).

**Figure 4.**
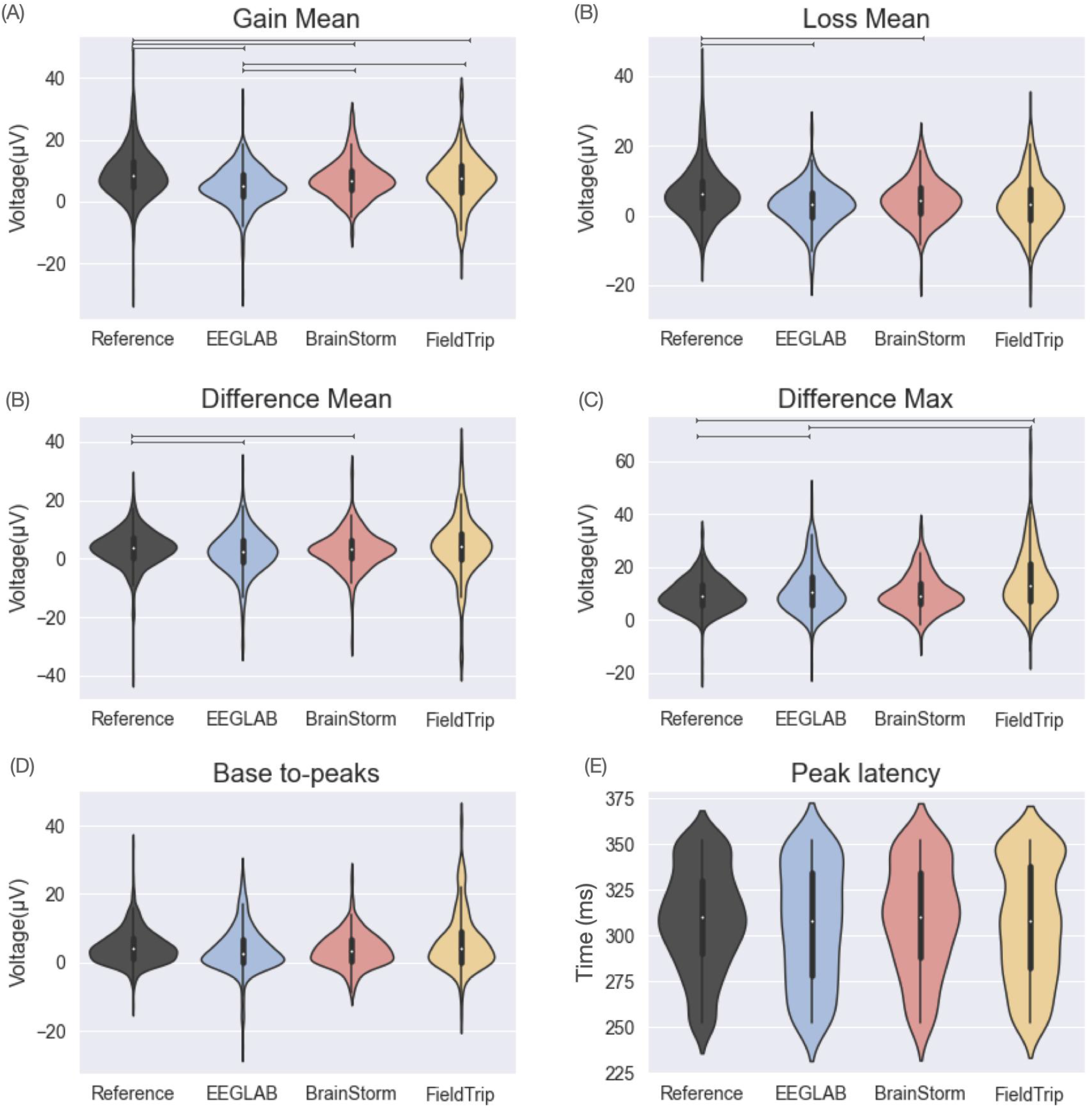
The violin plots showing the software distribution across subjects of the quantitative measures. A line between two violins denotes a statistical difference between their corresponding values.

**Figure 5.**
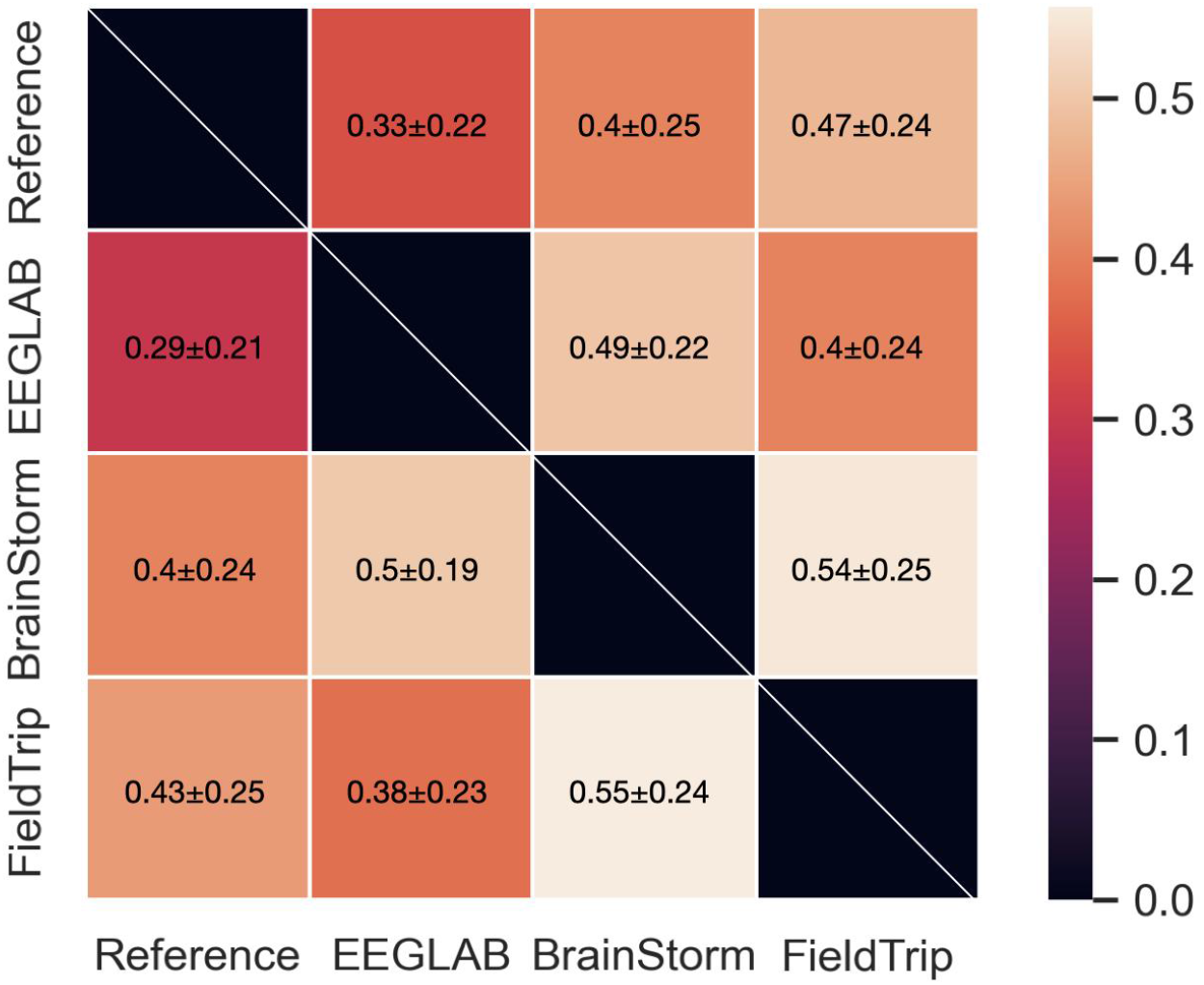
The similarity matrix between the conditional ERPs obtained by the different softwares. The upper triangular part of the matrix corresponds to the gain condition while the lower part corresponds to the loss condition.

## 3. Results

### 3.1. Reproduction of the main findings

In Figure 2, we illustrate the ERP waveforms at electrode FCz reflecting the reward positivity, obtained when running the preprocessing code of the reference paper, EEGLAB, BrainStorm and Fieldtrip. The grand averaged ERPs shown in Figure 2 are obtained after averaging all the clean epochs (kept after the artifactual trials removal step) of all subjects. For EEGLAB, the results illustrated in Figure 2 correspond to those obtained using the *clean_rawdata* function. Another automatic way incorporated in EEGLAB is the PREP pipeline (Bigdely-Shamlo et al. 2015) that uses four metrics to identify bad channels: extreme amplitudes (deviation criterion), lack of correlation with any other channel (correlation criterion), lack of predictability by other channels (predictability criterion), and unusual high frequency noise (noisiness criterion). The preprocessing results using the PREP pipeline can be found in the supplementary materials (see FIgure S1).

We observed a good degree of consistency between the ERP results published in the original paper and those reproduced using the script provided but excluding the ICA step (see Methods “Original preprocessing pipeline” for more details). The peak amplitudes of the conditional ERPs observed at 400 ms latency were slightly higher to those revealed when eye blinks were removed using ICA. Quantitatively, the same conclusions regarding the effect size related to the maximum and base to peak measures of the difference ERP, and the mean measure of the conditional ERPs were derived.

Changes in the amplitude of the two peaks were noticed for the conditional ERP generated by EEGLAB, Brainstorm and Fieldtrip, when compared to the reference results. A remarkable variability in the number of remained trials/subjects obtained after the preprocessing was observed between the software tools: the number of subjects with clean data was N=498 for the reference preprocessing, N=300 for EEGLAB, N=213 for Brainstorm code, and N=397 for FieldTrip code. This change in the number of participants is clearly reflected in the width of the confidence intervals (CI) which was larger for EEGLAB, BrainStorm and FieldTrip compared to the reference. Despite those differences, there was a good level of concordance between the conditional ERPs obtained in terms of the two peaks latencies seen respectively at 250 ms and 400 ms for all the software tools. In addition, the same waveform profile showing positive and negative deflections at specific times was visualized. One can also notice that, for all software tools, the gain waveform is above or at the same level as the loss waveform between 0 ms and 450 ms whereas the gain ERP passes under the loss ERP for the remaining time.

According to the grand averaged difference, the reward positivity peaked at a latency of 310 ms for the different softwares. While the peak voltage is approximately conserved for Brainstorm and FieldTrip, it is decreased after EEGLAB preprocessing (reaching 3 microvolts compared to 5 microVolts for the reference peak voltage).

Looking at the quantitative measures, results show good consistency between the reference and the software tools (Figure 3, table 1, table 2). More specifically, a large effect size (d>0.8) is obtained when looking at the maximum and base to peak measures of the difference ERP obtained by all the software tools (table 1, last column). For the mean peak metric, results of all software except EEGLAB present a medium effect size (d>0.5). Moreover, the conditional gain ERP elicited a mean peak measure with a large effect size (d>0.8) for all the software tools. However, the effect size of the mean peak related to the conditional loss ERP was only considered large for the reference and the FieldTrip results (d>0.8), but not for EEGLAB and Brainstorm (d<0.8). Overall the absolute voltage observed for the gain and loss ERPs separately were consistently smaller for all studied software packages compared to the reference paper.

**Table 1.** Mean, maximum and base to peak and the effect size of the reward positivity (the difference ERP) for the reference paper as well as the three studied software packages EEGLAB BrainStorm and FieldTrip. The reported mean, standard deviation and cohen’s d values were computed across subjects. This table is reproduced from Table 1 reported in (Williams et al. 2021).

|  |  | Mean [95% CI] | Standard deviation | Cohen's d [95% CI] |
| --- | --- | --- | --- | --- |
| **Original paper**<br>**(Williams et al. 2021)** | **Mean** | 3.70 [3.34 , 4.07 ] | 4.11 | 0.90 [0.77, 1.03] |
|  | **Maximum** | 7.82 [7.42 , 8.23 ] | 4.59 | 1.71 [1.56, 1.85] |
|  | **Base to peak** | 10.52 [10.12 , 10.91 ] | 4.49 | 2.34 [2.18, 2.50] |
| **Reference** | **Mean** | 3.45 [ 2.75 4.15] | 7.94 | 0.62 [0.48 0.75 ] |
|  | **Maximum** | 9.73 [ 9.17 10.28] | 6.31 | 2.18 [1.95 2.41] |
|  | **Base to peak** | 14 [13.34 14.64] | 7.36 | 2.69 [2.42 2.96] |
| **EEGLAB** | **Mean** | 2.36 [1.5 3.22] | 7.56 | 0.44 [ 0.27 0.61] |
|  | **Maximum** | 11.82 [10.8 12.8] | 8.8 | 1.9 [1.63 2.2] |
|  | **Base to peak** | 18.2 [17.03 19.42] | 10.4 | 2.46 [2.14 2.79] |
| **BrainStorm** | **Mean** | 3.25 [ 2.37 4.13] | 6.6 | 0.69 [ 0.49 0.91] |
|  | **Maximum** | 10.47 [9.53 11.42] | 7.11 | 2.08 [ 1.75 2.4] |
|  | **Base to peak** | 14.95 [13.9 16] | 7.8 | 2.70 [ 2.29 3.11] |
| **FieldTrip** | **Mean** | 4.28 [ 3.35 5.21] | 10.54 | 0.57 [0.44 0.71] |
|  | **Maximum** | 15.54 [14.5 16.5] | 11.58 | 1.9 [1.7 2.1] |
|  | **Base to peak** | 23.47 [ 22.14 24.8] | 15.09510 | 2.2 [ 1.97 2.43] |

**Table 2.** Effect size of the gain and loss conditional ERP, using the meak peak measure, for all software. The reported mean, standard deviation and cohen’s d values were computed across subjects. This table is reproduced from Table 2 reported in (Williams et al. 2021).

|  |  | Mean [95% CI] | Standard deviation | Cohen's d [95% CI] |
| --- | --- | --- | --- | --- |
| **Original paper**<br>**(Williams et al. 2021)** | **Gain** | 8.02 [7.54 , 8.49 ] | 5.38 | 1.49 [1.35, 1.63] |
|  | **Loss** | 4.96 [4.53 , 5.38 ] | 4.87 | 1.02 [0.89, 1.15] |
| **Reference** | **Gain** | 9.05 [8.37 9.7] | 7.8 | 1.64 [1.45 1.83] |
|  | **Loss** | 6.80[6.11 7.5] | 7.84 | 1.23 [1.06 1.4 ] |
| **EEGLAB** | **Gain** | 4.91 [4.19 5.63] | 6.3 | 1.09 [0.89 1.3] |
|  | **Loss** | 2.87 [2.2 3.5] | 5.8 | 0.69 [0.51 0.87] |
| **BrainStorm** | **Gain** | 7.31 [6.5 8.14] | 6.2 | 1.67 [1.38 1.96] |
|  | **Loss** | 4.59 [3.79 5.39] | 5.9 | 1.08 [0.84 1.32] |
| **FieldTrip** | **Gain** | 7.21 [6.5 7.9] | 8.15 | 1.25 [1.08 1.42] |
|  | **Loss** | 3.61 [2.9 4.2] | 7.63 | 0.67 [0.53 0.81] |

### 3.2. Comparison across software

A significant statistical difference was observed between the reference and all the software toolboxes in terms of the mean peak amplitude of the gain ERP (Figure 4). Significant statistical differences between the mean peak voltages of the loss and the difference ERP of the reference and those of EEGLAB and Brainstorm softwares were also observed. FieldTrip shows statistically different maximum peak of the difference ERP when compared to the reference. Regarding the base-to-peak and the peak latency measures, no statistical differences were found between softwares.

Figure 5 illustrates the similarity matrix between the conditional ERPs generated by the different softwares when taking into account all the EEG channels. For each participant, the similarity between the preprocessed ERPs obtained from two different softwares was calculated using pearson’s correlation averaged across all channels (see materials and methods). Between the three Matlab toolboxes, FieldTrip reached the highest similarity for both conditions (0.47±0.24 for the gain condition; 0.43±0.25 for the loss condition), followed by Brainstorm (0.4 ±0.25 for the gain condition and 0.4 ±0.24 for the loss condition) then EEGLAB (0.33 ±0.22 for the gain condition and 0.29 ±0.21 for the loss condition). The highest similarity is observed between Brainstorm and FieldTrip (0.54 ±0.25 for the gain condition, and 0.55 ±0.24 for the loss condition).

In addition, we investigated where and when the ERPs were statistically different by plotting the thresholded statistical map (time x channels). To do this, we quantified the statistical difference between the subjects’ distribution of ERPs obtained from two softwares, at each millisecond and each channel using the Wilcoxon test (see materials and methods). In line with the previous findings, Figure 6 highlights that EEGLAB shows the highest statistical differences compared to the reference results. One can also remark that EEGLAB statistical differences are distributed along the time axis starting from 100 ms to 1000 ms after the stimulus. The major statistical differences between FieldTrip and the reference results are revealed between 300 ms and 700 ms, at all the EEG channels adequately. While both the gain and loss conditions show important differences for all three software packages compared to the reference paper, there is only limited areas of significant differences in the reward positivity (i.e. ERP difference between gain and loss conditions).

**Figure 6.**
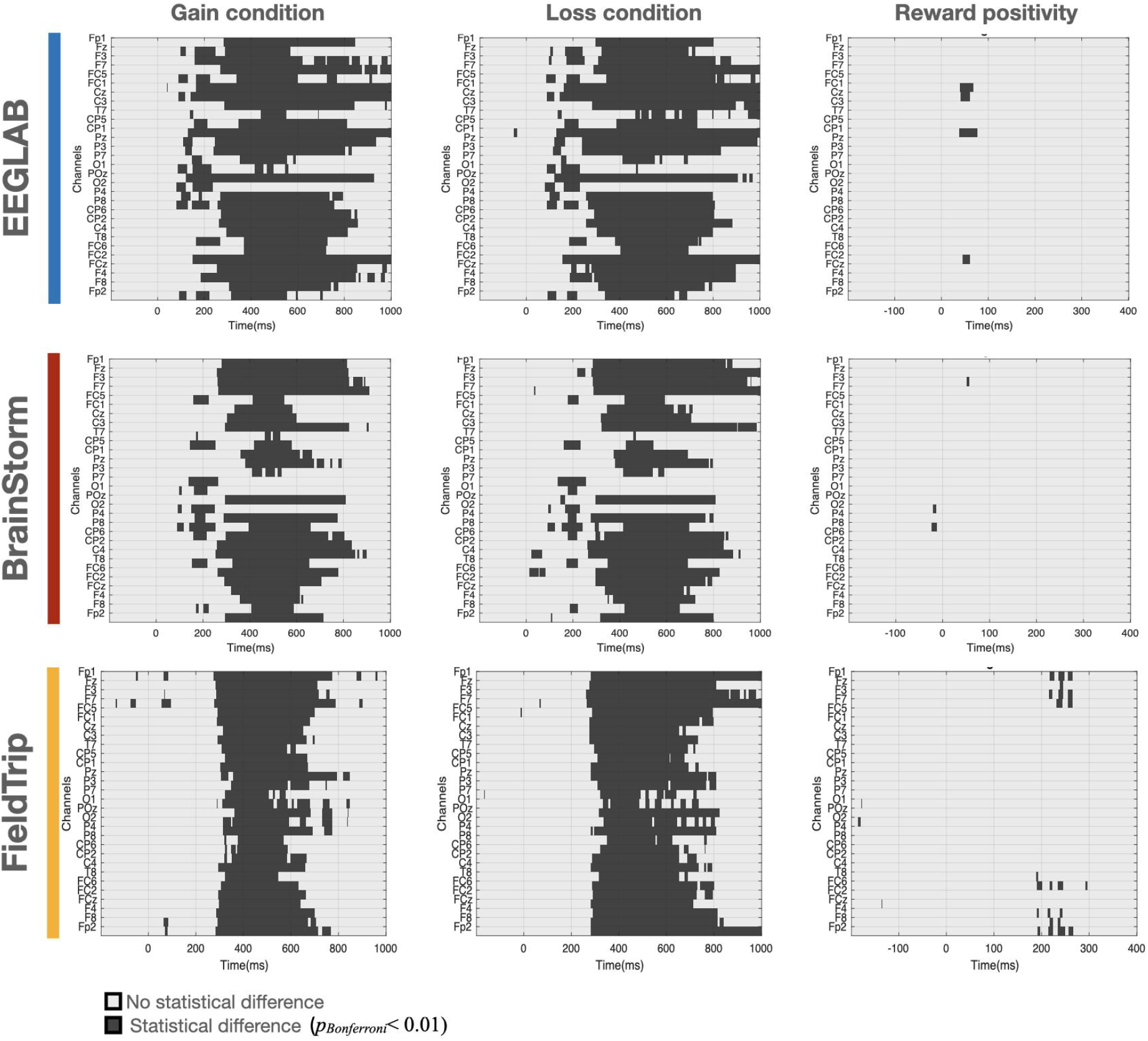
The statistical maps showing the differences between the results of each software tool and the reference at each millisecond and channel.

## 4. Discussion

A large range of techniques and tools are now available to process a single EEG dataset. This high analytical flexibility, reflected by the large number of choices made during the data preprocessing and analysis workflow, can be problematic as it can yield variability in research outcomes. Therefore, it is important to particularly understand the impact of the preprocessing methods, software package, software version and even the operating system on the reproducibility of the final research outcome of a study.

Here, we were interested in exploring the impact of the preprocessing software on the ERP derived from EEG data of 500 participants performing a simple gambling task as originally published by (Williams et al. 2021). The degree of agreement across software packages was good in terms of peak latencies and the general profile of ERP waveforms. In addition, the majority of the tested softwares obtained similar effect size estimates related to specific ERP features. However, remaining variability was also observed between software packages. This variability was reflected by the number of clean trials kept to compute the grand averaged ERPs, the peak voltages, the width of the confidence interval, and the statistical differences at particular channels and time instants (due to differences in absolute voltage values). Among the tested softwares used to reproduce the same preprocessing pipeline published by (Williams et al. 2021), EEGLAB seems to generate results with the lowest similarity when compared to the original ones while Brainstorm generates results with the highest similarity. However, it is noteworthy to pinpoint that the objective of the current study is not to favor any software tool over another or to recommend the ‘best’ preprocessing tool, but rather, to illuminate and quantify differences that can be generated by different softwares on the same database. The variations observed can be related to several factors that are implicated in the preprocessing steps applied in each software. Results are discussed hereafter.

### 4.1. Influencing factors

In this study, our objective was to re-analyze the same data originally published and preprocessed in (Williams et al. 2021) using EEGLAB, Brainstorm and FieldTrip following the same original preprocessing workflow. Our intent was to fully automate all the preprocessing operations avoiding any manual intervention as much as practicable. Computationally, the workflow in each software was designed as a sequence of steps that are combined so that the intermediate outputs from one step directly feed as inputs into the next step. Notably, while all the tested software toolboxes were purportedly replicating the same preprocessing steps, it was often impossible to exactly adapt the same methods and choice parameters used in the reference paper due to software implementation and configuration choices..

For instance, the band-pass filter cannot be configured in EEGLAB and Brainstorm to have the same type (Butterworth) and order used in the original paper. In addition, the gradient criteria adapted by the original study to detect the bad epochs is not supported by any of the tested software. More crucially, the method performed to detect the bad channels contributed to the major disparities seen across packages in terms of the number of trials and subjects used for ERP computation. This is because the bad channels interpolated by the original pipeline were identified in a way to have a low trial rejection rate (see materials and methods section for more details). To do so, data were segmented into trials twice wherein the first pass was employed to identify the noisy channels, and the second pass was to process data for analysis. This particular process has not only identified very specific channels to be interpolated, but also it leads to a high number of ‘good’ trials that will be used ultimately to reconstruct the ERPs. In EEGLAB, Brainstorm and FieldTrip, the automatic detection of bad channels was carried out using their designed functions based on different criteria. This can explain the considerable difference between softwares in the number of trials and subjects remaining after the data preprocessing.

To better investigate whether the results discrepancies are mainly dependent on the number of subjects showing clean data, we compared the results of ERPs computed using data corresponding to the common subjects remaining after the preprocessing across the reference, EEGLAB, Brainstorm and FieldTrip (see Figure S2 in supplementary materials). Still, results show variability in the ERP waveforms and related characteristics, which means that other influencing factors are contributing to the observed variability. We also tried to regulate the parameters selected in the bad trial identification step to increase the number of clean trials kept by each software. Thus, instead of the 100 microVolts min-max criterion, we tested the variability of results between softwares when using 200 microVolts min-max criterion (see Figure S3, Table S1, Table S3 in supplementary materials). Compared to the results obtained using the 100 microVolts min-max criterion, higher similarities were markedly obtained for all the softwares. Related to all the ERP features (mean, max, base-to-peak and peak latency), the three softwares succeeded to validate the effect size estimates obtained by the reference results. The number of subjects showing clean data was 426 after running EEGLAB code, 316 after Brainstorm code, and 497 after FieldTrip code. Remarkably, EEGLAB still showed more statistical differences with the reference results, compared to Brainstorm and Fieldtrip. In contrast, Brainstorm results revealed the highest similarity to the reference results even when only 316 subjects were kept. This highlights the need for a multi-stage assessment of software differences, to examine which steps made the major difference in study outcomes, and which steps were of less concern.

### 4.2. Reproducibility in the neuroimaging field

The question of reproducibility and replicability is considerably gaining attention in the scientific community (Nosek et al. 2022; Fidler and Wilcox 2018; Munafò et al. 2020). In the neuroimaging field, a recent study addressed the issue of analytical flexibility in fMRI research and its effects on the associated conclusions (Botvinik-Nezer et al. 2020). Using the same data, variability in results was reported in testing nine hypotheses across seventy independent teams. Inspired by this study, two recent initiatives have been made to test the effect of diversity of analysis pipelines and teams on EEG results. The ‘EEGManyPipelines’ (Algermissen, Yang, and Busch 2021) project and EEGManyLabs (Pavlov et al. 2021) were recently launched to involve many independent teams in analyzing the same data and testing a set of predefined hypotheses. Multiple EEG studies have also demonstrated that a study outcomes are contingent on subjective decisions and factors selected in the EEG analysis, such as the EEG electrode density (Song et al. 2015; Sohrabpour et al. 2015; Lantz et al. 2003; Allouch et al. 2022), the preprocessing methods and parameters (Šoškić et al. 2022; Barban et al. 2021; Robbins et al. 2020; Clayson et al. 2021), the number of trials (Boudewyn et al. 2018) and the specific parameters related to the EEG connectivity analysis (Allouch et al. 2022; Hassan et al. 2014). The variability in the software used in EEG analysis was tackled in a recent review that addresses the question of reproducibility and consistency of ERP studies, mainly focusing on the N400 component (Šoškić et al. 2021). In a sample of 132 ERP papers, (Šoškić et al. 2021) reveals that the number of softwares used to perform the EEG analysis stages (from the presentation of stimulus to the statistical assessment) ranged from 8 to 17 options, and that such methodological decision can induce substantial variability in the reported results, ultimately hindering research replicability. Using fMRI, many studies have quantified the impacts of the analysis software (Bowring, Maumet, and Nichols 2019; Li et al. 2021), the software version (Gronenschild et al. 2012) and the operating system (Glatard et al. 2015; Gronenschild et al. 2012) on results conducted on a single dataset. In the current study, we focused on examining whether it is possible to reproduce the same ERP results after preprocessing data with different software tools. To the best of our knowledge, the effect of the preprocessing software on the same EEG dataset has never been studied before. This current study has not only provided a validation of EEGLAB, Brainstorm and FieldTrip but also it contributed to better understand the possible discrepancies in results generated by different EEG studies.

### 4.3. Methodological considerations

In this work, we attempted to reproduce? using different softwares? the same preprocessing pipeline initially proposed by (Williams et al. 2021). This was carefully done by conserving, as much as possible, the same steps along with their related parameters (cut-off filters, baseline duration, reference electrodes..etc) and order. Nevertheless, the derived signals and results might be also sensitive to other factors that were not investigated in this study. For instance, the parameters used to detect the artifactual channels were set to the default or to the most commonly used values as recommended by each toolbox (such as the window length in which signals are completely flat, the correlation with neighbors threshold and other criteria). An interesting future prospect would be testing the consistency of results when varying these factors.

To validate the hypotheses supported by the reference paper exploring the same dataset, we compared the results generated by each software to those originally published by (Williams et al. 2021). The cross-software discrepancies were also quantified. Another issue that may be of great interest to be investigated is to evaluate the feasibility of each preprocessing tool in generating reliable signals with good data quality. This could be done by comparing results to ground-truth data generated ideally by a computational model of electrophysiological signals such as neural-mass models (Bensaid et al. 2019) or multivariate autoregressive models (Anzolin et al. 2019; Haufe and Ewald 2016)

In order to conduct the comparative analysis between the results generated by the different softwares, we used several quantification metrics to measure the consistency/discrepancy of ERP waveforms and their related characteristics. We mainly relied on ERP as the main objective of this work was to reproduce and validate the results published by (Williams et al. 2021) studying the reward positivity. However, it is commonly known that ERP strategy is based on across-trial averaging which increases the signal-to-noise ratio, and blinds a considerable part of the elicited brain activity. Thus, we are aware that the consistency of results may greatly differ if the analysis was performed on the continuous preprocessed EEG instead of ERPs computed after averaging a large number of epochs. In addition, considering smaller sample sizes could also lead to higher levels of cross-software variability as previous literature has outlined how variability induced by different pipelines decreases with higher signal-to-noise ratio (e.g. see (Li et al. 2021) for an example with resting rate fMRI of various acquisition durations).

Among the available preprocessing tools used in EEG studies, we selected three of the most commonly used softwares. In each software, we tried to follow, as much as possible, the same preprocessing workflow of (Williams et al. 2021) using the provided software functions. This led us to exclude other interesting packages that conduct a fully automatic preprocessing such as automagic (Pedroni, Bahreini, and Langer 2019), the Harvard Automated Preprocessing Pipeline for EEG (HAPPE) (Gabard-Durnam et al. 2018) and the Batch Electroencephalography Automated Processing Platform (BEAPP) (Levin et al. 2018) toolboxes. In other words, our inability to control or modify the inclusion and the order of the various preprocessing steps impedes these toolboxes to respect the same preprocessing pipeline we were trying to reproduce. Besides Matlab, it would be interesting to investigate and systematically quantify the differences of results generated by the MNE-python package (Gramfort et al. 2013). In addition, it is unclear how findings reported in this paper would generalize to other datasets or experimental paradigms. Therefore, it would be interesting to evaluate the fluctuations of results on other datasets and tasks covering further preprocessing pipelines and steps.

## 5. Conclusions

This study sheds light on how the software tool used to preprocess EEG signals impacts the analysis results and conclusions. EEGLAB, Brainstorm and FieldTrip were used to reproduce the same preprocessing pipeline of a published EEG study performed on 500 participants. While the three softwares succeeded to infer the same conclusion of the original publication regarding the effect size estimates related to the ERP features derived, we observed significant differences in terms of the observed absolute voltage between EEGLAB, Brainstorm and Fieldtrip results, as well as between each of the softwares and the original results. To better understand the variability induced by the software tool, further comparative studies should be conducted to examine the effects on the continuous EEG signals instead of ERP signals. In addition, more in-depth analysis is recommended in order to identify the critical steps and factors that lead the most to the variability observed.

## 6. Acknowledgments

This work was supported by the Institute of Clinical Neuroscience of Rennes (Projects named EEGNET3). Authors would like to thank Campus France, Programme Hubert Curien CEDRE (PROJET N° 42257YA) and the Lebanese Association for Scientific Research (LASER) for their support.

## References

Algermissen, J., Y. F. Yang, and N. A. Busch. 2021. “EEGManyPipelines: Mapping the Diversity of EEG Analysis Pipelines and Their Impact on Results.” https://repository.ubn.ru.nl/handle/2066/241382.

Allouch, Sahar, Maxime Yochum, Aya Kabbara, Joan Duprez, Mohamad Khalil, Fabrice Wendling, Mahmoud Hassan, and Julien Modolo. 2022. “Mean-Field Modeling of Brain-Scale Dynamics for the Evaluation of EEG Source-Space Networks.” Brain Topography 35 (1): 54–65.

Anzolin, Alessandra, Paolo Presti, Frederik Van De Steen, Laura Astolfi, Stefan Haufe, and Daniele Marinazzo. 2019. “Quantifying the Effect of Demixing Approaches on Directed Connectivity Estimated Between Reconstructed EEG Sources.” Brain Topography. https://doi.org/10.1007/s10548-019-00705-z.

Barban, Federico, Michela Chiappalone, Gaia Bonassi, Dante Mantini, and Marianna Semprini. 2021. “Yet Another Artefact Rejection Study: An Exploration of Cleaning Methods for Biological and Neuromodulatory Noise.” Journal of Neural Engineering 18 (4). https://doi.org/10.1088/1741-2552/ac01fe.

Bensaid, Siouar, Julien Modolo, Isabelle Merlet, Fabrice Wendling, and Pascal Benquet. 2019. “COALIA: A Computational Model of Human EEG for Consciousness Research.” Frontiers in Systems Neuroscience 13 (November): 59.

Bigdely-Shamlo, Nima, Tim Mullen, Christian Kothe, Kyung-Min Su, and Kay A. Robbins. 2015. “The PREP Pipeline: Standardized Preprocessing for Large-Scale EEG Analysis.” Frontiers in Neuroinformatics 9 (June): 16.

Botvinik-Nezer, Rotem, Felix Holzmeister, Colin F. Camerer, Anna Dreber, Juergen Huber, Magnus Johannesson, Michael Kirchler, et al. 2020. “Variability in the Analysis of a Single Neuroimaging Dataset by Many Teams.” Nature 582 (7810): 84–88.

Boudewyn, Megan A., Steven J. Luck, Jaclyn L. Farrens, and Emily S. Kappenman. 2018. “How Many Trials Does It Take to Get a Significant ERP Effect? It Depends.” Psychophysiology 55 (6): e13049.

Bowring, Alexander, Camille Maumet, and Thomas E. Nichols. 2019. “Exploring the Impact of Analysis Software on Task fMRI Results.” Human Brain Mapping 40 (11): 3362–84.

Button, Katherine S., John P. A. Ioannidis, Claire Mokrysz, Brian A. Nosek, Jonathan Flint, Emma S. J. Robinson, and Marcus R. Munafò. 2013. “Power Failure: Why Small Sample Size Undermines the Reliability of Neuroscience.” Nature Reviews. Neuroscience 14 (5): 365–76.

Clayson, Peter E., Scott A. Baldwin, Harold A. Rocha, and Michael J. Larson. 2021. “The Data-Processing Multiverse of Event-Related Potentials (ERPs): A Roadmap for the Optimization and Standardization of ERP Processing and Reduction Pipelines.” NeuroImage 245 (December): 118712.

Croft, Rodney J., Jody S. Chandler, Robert J. Barry, Nicholas R. Cooper, and Adam R. Clarke. 2005. “EOG Correction: A Comparison of Four Methods.” Psychophysiology 42 (1): 16–24.

Delorme, Arnaud, and Scott Makeig. 2004. “EEGLAB: An Open Source Toolbox for Analysis of Single-Trial EEG Dynamics Including Independent Component Analysis.” Journal of Neuroscience Methods 134 (1): 9–21.

Fidler, Fiona, and John Wilcox. 2018. “Reproducibility of Scientific Results.” https://stanford.library.sydney.edu.au/archives/win2019/entries/scientific-reproducibility/.

Gabard-Durnam, Laurel J., Adriana S. Mendez Leal, Carol L. Wilkinson, and April R. Levin. 2018. “The Harvard Automated Processing Pipeline for Electroencephalography (HAPPE): Standardized Processing Software for Developmental and High-Artifact Data.” Frontiers in Neuroscience 12. https://doi.org/10.3389/fnins.2018.00097.

Glatard, Tristan, Lindsay B. Lewis, Rafael Ferreira da Silva, Reza Adalat, Natacha Beck, Claude Lepage, Pierre Rioux, et al. 2015. “Reproducibility of Neuroimaging Analyses across Operating Systems.” Frontiers in Neuroinformatics 9 (April): 12.

Gramfort, Alexandre, Martin Luessi, Eric Larson, Denis A. Engemann, Daniel Strohmeier, Christian Brodbeck, Roman Goj, et al. 2013. “MEG and EEG Data Analysis with MNE-Python.” Frontiers in Neuroscience 7 (December): 267.

Gramfort, Alexandre, Martin Luessi, Eric Larson, Denis A. Engemann, Daniel Strohmeier, Christian Brodbeck, Lauri Parkkonen, and Matti S. Hämäläinen. 2014. “MNE Software for Processing MEG and EEG Data.” NeuroImage 86 (February): 446–60.

Gronenschild, Ed H. B. M., Petra Habets, Heidi I. L. Jacobs, Ron Mengelers, Nico Rozendaal, Jim van Os, and Machteld Marcelis. 2012. “The Effects of FreeSurfer Version, Workstation Type, and Macintosh Operating System Version on Anatomical Volume and Cortical Thickness Measurements.” PloS One 7 (6): e38234.

Hassan, Mahmoud, Olivier Dufor, Isabelle Merlet, Claude Berrou, and Fabrice Wendling. 2014. “EEG Source Connectivity Analysis: From Dense Array Recordings to Brain Networks.” PloS One 9 (8): e105041.

Haufe, Stefan, and Arne Ewald. 2016. “A Simulation Framework for Benchmarking EEG-Based Brain Connectivity Estimation Methodologies.” Brain Topography 32 (4): 625–42.

Ioannidis, John P. A. 2005. “Why Most Published Research Findings Are False.” PLoS Medicine 2 (8): e124.

Islam, Md Kafiul, Amir Rastegarnia, and Zhi Yang. 2016. “Methods for Artifact Detection and Removal from Scalp EEG: A Review.” Neurophysiologie Clinique = Clinical Neurophysiology 46 (4-5): 287–305.

Lantz, Goran, R. Grave de Peralta, L. Spinelli, M. Seeck, and C. M. Michel. 2003. “Epileptic Source Localization with High Density EEG: How Many Electrodes Are Needed?” Clinical Neurophysiology: Official Journal of the International Federation of Clinical Neurophysiology 114 (1): 63–69.

Levin, April R., Adriana S. Méndez Leal, Laurel J. Gabard-Durnam, and Heather M. O’Leary. 2018. “BEAPP: The Batch Electroencephalography Automated Processing Platform.” Frontiers in Neuroscience 12 (August): 513.

Li, Xinhui, Lei Ai, Steve Giavasis, Hecheng Jin, Eric Feczko, Ting Xu, Jon Clucas, et al. 2021. “Moving Beyond Processing and Analysis-Related Variation in Neuroscience.” bioRxiv. https://doi.org/10.1101/2021.12.01.470790.

Lopes da Silva, Fernando. 2013. “EEG and MEG: Relevance to Neuroscience.” Neuron 80 (5): 1112–28.

Matlab. 2018. “9.5.0.944444 (R2018b).” Natick, Massachusetts: The MathWorks Inc.

Munafò, Marcus R., Christopher D. Chambers, Alexandra M. Collins, Laura Fortunato, and Malcolm R. Macleod. 2020. “Research Culture and Reproducibility.” Trends in Cognitive Sciences 24 (2): 91–93.

Nosek, Brian A., Tom E. Hardwicke, Hannah Moshontz, Aurélien Allard, Katherine S. Corker, Anna Dreber, Fiona Fidler, et al. 2022. “Replicability, Robustness, and Reproducibility in Psychological Science.” Annual Review of Psychology 73 (January): 719–48.

Oostenveld, Robert, Pascal Fries, Eric Maris, and Jan-Mathijs Schoffelen. 2011. “FieldTrip: Open Source Software for Advanced Analysis of MEG, EEG, and Invasive Electrophysiological Data.” Computational Intelligence and Neuroscience 2011: 156869.

Pavlov, Adamian, Appelhoff, and Arvaneh. 2021. “# EEGManyLabs: Investigating the Replicability of Influential EEG Experiments.” Cortex; a Journal Devoted to the Study of the Nervous System and Behavior. https://www.sciencedirect.com/science/article/pii/S0010945221001106.

Pedroni, Andreas, Amirreza Bahreini, and Nicolas Langer. 2019. “Automagic: Standardized Preprocessing of Big EEG Data.” NeuroImage 200 (October): 460–73.

Picton, T. W., O. G. Lins, and M. Scherg. 1995. “The Recording and Analysis of Event-Related Potentials.” Handbook of Neuropsychology. https://www.researchgate.net/profile/Terence-Picton/publication/247966238_The_recording_and_analysis_of_event-related_potentials/links/552e75a20cf2d495071844ee/The-recording-and-analysis-of-event-related-potentials.pdf.

Proudfit, Greg Hajcak. 2015. “The Reward Positivity: From Basic Research on Reward to a Biomarker for Depression.” Psychophysiology 52 (4): 449–59.

Ranjan, Rakesh, Bikash Chandra Sahana, and Ashish Kumar Bhandari. 2021. “Ocular Artifact Elimination from Electroencephalography Signals: A Systematic Review.” Biocybernetics and Biomedical Engineering 41 (3): 960–96.

R Core Team. 2020. R: A Language and Environment for Statistical Computing. Vienna, Austria: R Foundation for Statistical Computing. https://www.R-project.org/.

Robbins, Kay A., Jonathan Touryan, Tim Mullen, Christian Kothe, and Nima Bigdely-Shamlo. 2020. “How Sensitive Are EEG Results to Preprocessing Methods: A Benchmarking Study.” IEEE Transactions on Neural Systems and Rehabilitation Engineering: A Publication of the IEEE Engineering in Medicine and Biology Society 28 (5): 1081–90.

Sambrook, Thomas D., and Jeremy Goslin. 2015. “A Neural Reward Prediction Error Revealed by a Meta-Analysis of ERPs Using Great Grand Averages.” Psychological Bulletin 141 (1): 213–35.

Sohrabpour, Abbas, Yunfeng Lu, Pongkiat Kankirawatana, Jeffrey Blount, Hyunmi Kim, and Bin He. 2015. “Effect of EEG Electrode Number on Epileptic Source Localization in Pediatric Patients.” Clinical Neurophysiology: Official Journal of the International Federation of Clinical Neurophysiology 126: 472–80.

Song, J., C. Davey, C. Poulsen, P. Luu, S. Turovets, E. Anderson, K. Li, and D. Tucker. 2015. “EEG Source Localization: Sensor Density and Head Surface Coverage.” Journal of Neuroscience Methods 256: 9–21.

Šoškić, Anđela, Vojislav Jovanović, Suzy J. Styles, Emily S. Kappenman, and Vanja Ković. 2021. “How to Do Better N400 Studies: Reproducibility, Consistency and Adherence to Research Standards in the Existing Literature.” Neuropsychology Review, August. https://doi.org/10.1007/s11065-021-09513-4.

Šoškić, Anđela, Suzy J. Styles, Emily S. Kappenman, and Vanja Kovic. 2022. “Garden of Forking Paths in ERP Research –Effects of Varying Pre-Processing and Analysis Steps in an N400 Experiment.” https://doi.org/10.31234/osf.io/8rjah.

Tadel, François, Sylvain Baillet, John C. Mosher, Dimitrios Pantazis, and Richard M. Leahy. 2011. “Brainstorm: A User-Friendly Application for MEG/EEG Analysis.” Computational Intelligence and Neuroscience 2011 (April): 879716.

Urigüen, Jose Antonio, and Begoña Garcia-Zapirain. 2015. “EEG Artifact Removal—state-of-the-Art and Guidelines.” Journal of Neural Engineering 12 (3): 031001.

Waskom, Michael. 2021. “Seaborn: Statistical Data Visualization.” Journal of Open Source Software 6 (60): 3021.

Williams, Chad C., Thomas D. Ferguson, Cameron D. Hassall, Wande Abimbola, and Olave E. Krigolson. 2021. “The ERP, Frequency, and Time-Frequency Correlates of Feedback Processing: Insights from a Large Sample Study.” Psychophysiology 58 (2): e13722.

