## supplementary materials for "Successful reproduction of a large EEG study across software packages"

A

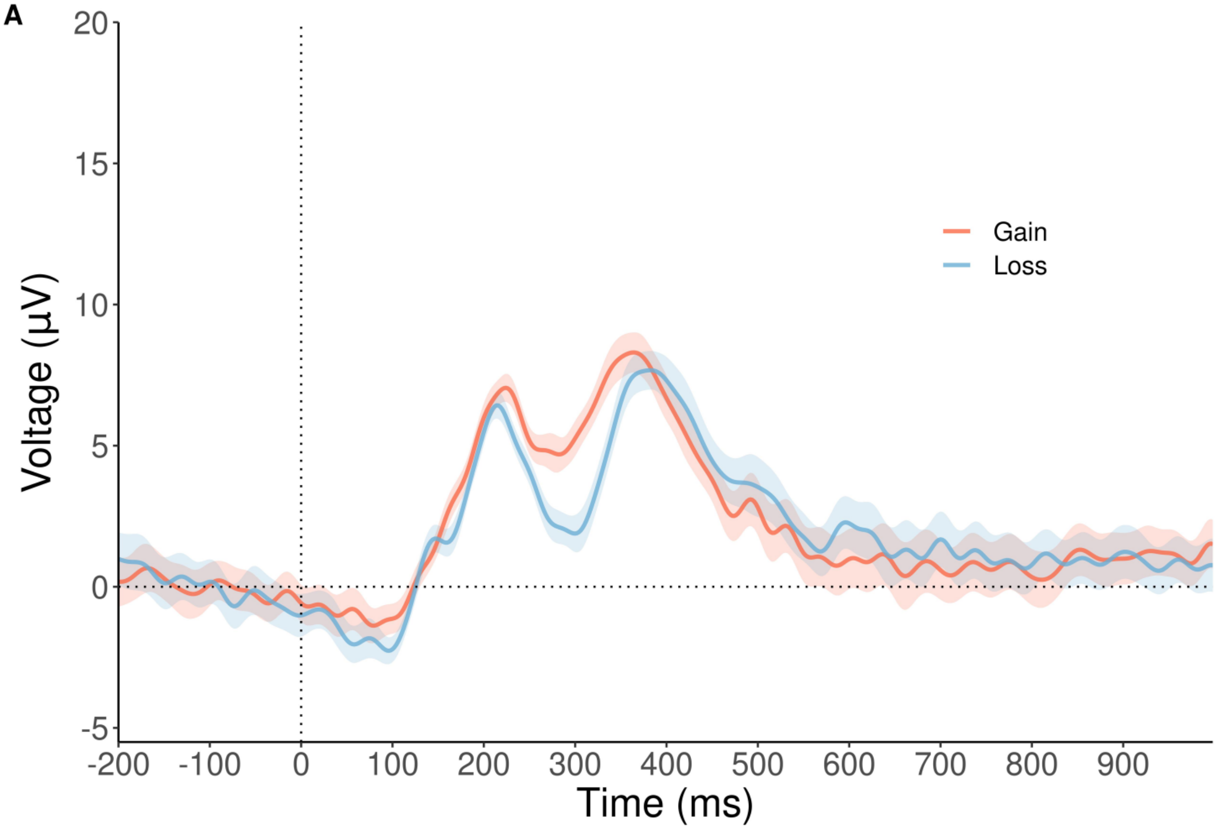

Gain
Loss
Voltage (µV)
20
15
10
5
0
-5
-200
-100
0
100
200
300
400
500
600
700
800
900
Time (ms)


B

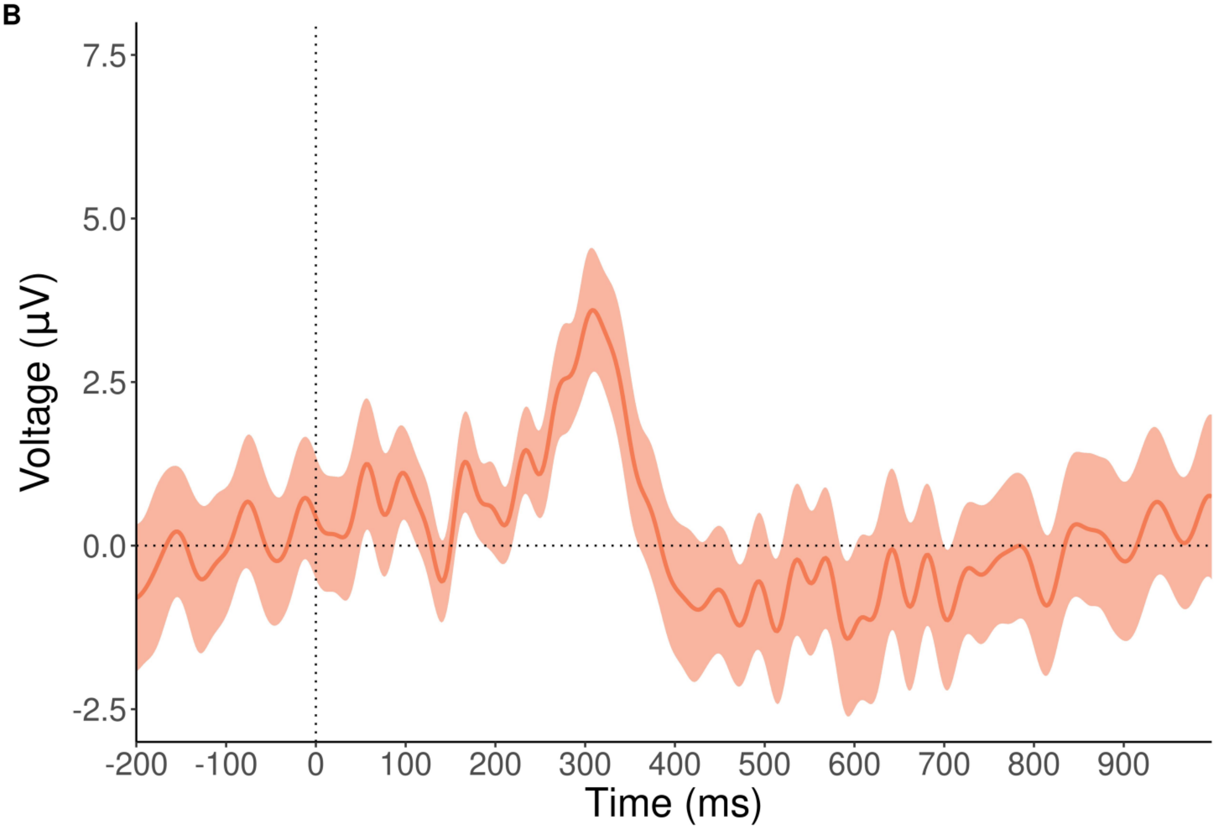

Voltage (µV)
7.5
5.0
2.5
0.0
-2.5
-200
-100
0
100
200
300
400
500
600
700
800
900
Time (ms)

Figure S1. ERP waveforms obtained using EEGLAB with PREP pipeline used to detect the bad channels. A) conditional ERPs for gain and loss condition, B) difference ERP between gain and loss

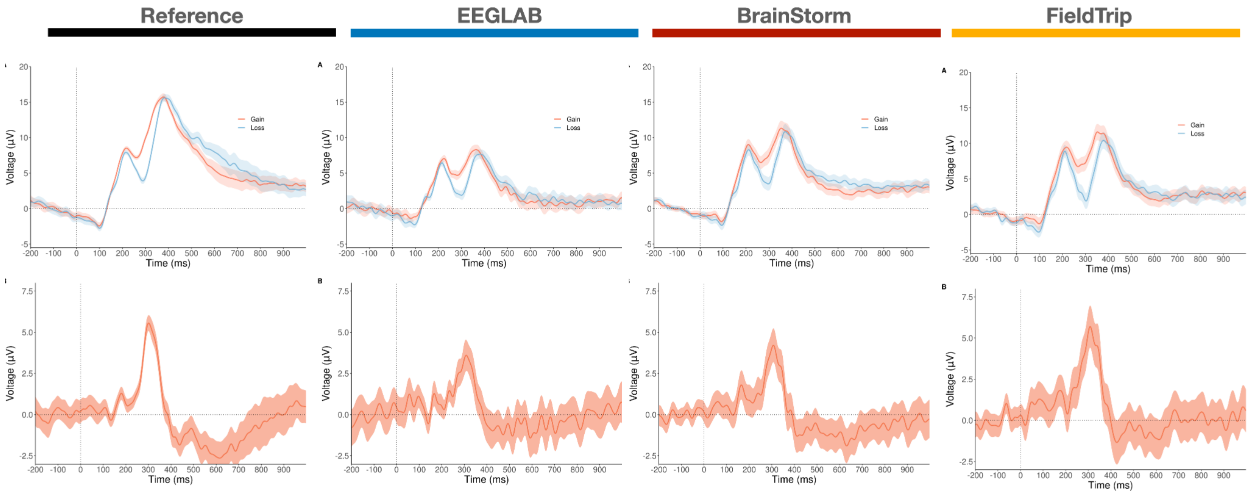


Figure S2. ERP waveforms obtained using EEGLAB with PREP pipeline used to detect the bad channels. A) conditional ERPs for gain and loss condition, B) difference ERP between gain and loss

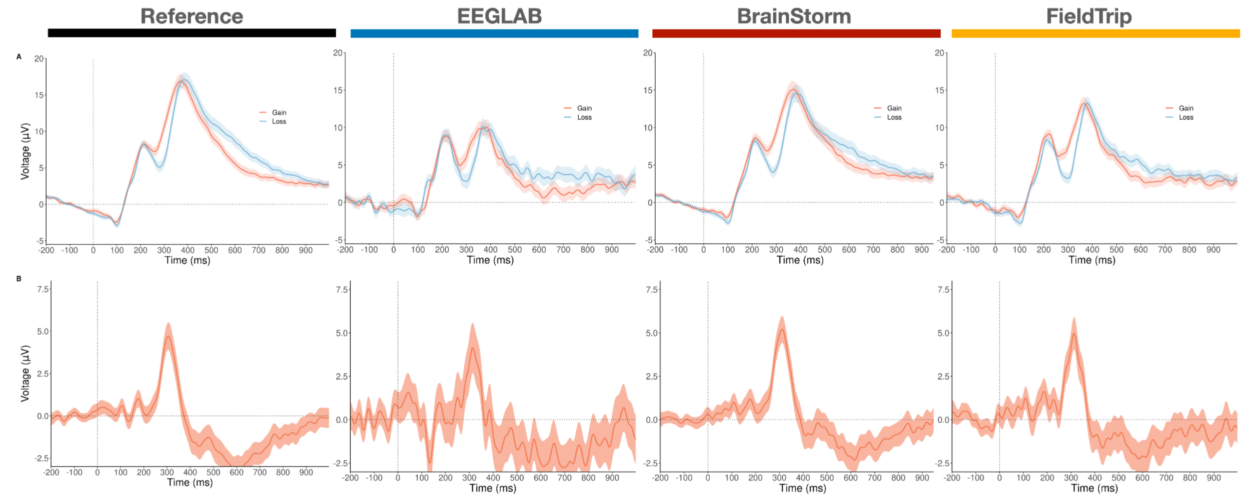

Figure S3. ERP waveforms at electrode FCz illustrating the reward positivity after preprocessing by the reference code, EEGLAB, Brainstorm and FieldTrip with rejecting trials exceeding 200 microVolts min-max. (a) grand averaged conditional waveforms (ERP averaged across all subjects) with 95% confidence intervals, (b) grand averaged difference waveform with 95% confidence intervals.

| | | Mean [95% CI] | Standard deviation | Cohen's d [95% CI] |
|---|---|---|---|---|
| **Original paper (Williams et al. 2021)** | **Mean** | 3.70 [3.34 , 4.07 ] | 4.11 | 0.90 [0.77, 1.03] |
| | **Maximum** | 7.82 [7.42 , 8.23 ] | 4.59 | 1.71 [1.56, 1.85] |
| | **Base to peak** | 10.52 [10.12 , 10.91 ] | 4.49 | 2.34 [2.18, 2.50] |
| **Reference** | **Mean** | 3.45 [ 2.75 4.15] | 7.94 | 0.62 [0.48 0.75 ] |
| | **Maximum** | 9.73 [ 9.17 10.28] | 6.31 | 2.18 [1.95 2.41] |
| | **Base to peak** | 14 [13.34 14.64] | 7.36 | 2.69 [2.42 2.96] |
| **EEGLAB** | **Mean** | 3.21[2.64 3.78] | 6 | 0.76 [0.61 0.9 ] |
| | **Maximum** | 10.41 [9.64 11.18] | 8.1 | 1.82 [1.6 2] |
| | **Base to peak** | 15.2 [14.31 16.19] | 9.9 | 2.18 [1.93 2.43] |
| **BrainStorm** | **Mean** | 3.84 [3.19 4.49] | 6.24 | 0.8 [0.69 1.04] |
| | **Maximum** | 9.74 [9.05 10.43] | 6.66 | 2.07 [1.81 2.32] |
| | **Base to peak** | 13.38 [12.6 14.13] | 7.22 | 2.62 [2.31 2.93] |
| **FieldTrip** | **Mean** | 3.4 [2.66 4.15] | 8.46 | 0.56 [0.43 0.7] |
| | **Maximum** | 12.22 [11.38 13] | 9.5 | 1.81 [1.61 2.01] |
| | **Base to peak** | 17.7 [16.79 18.79] | 11.3 | 2.22 [1.9 2.45] |

Table S1. Mean, maximum and base to peak and the effect size of the reward positivity (the difference ERP) for the reference paper as well as the three studied software packages EEGLAB BrainStorm and FieldTrip, with rejecting trials exceeding 200 microVolts min-max. The reported mean, standard deviation and cohen's d values were computed across subjects.

| | | Mean [95% CI] | Standard deviation | Cohen’s d [95% CI] |
|---|---|---|---|---|
| **Original paper (Williams et al. 2021)** | **Gain** | 8.02 [7.54 , 8.49 ] | 5.38 | 1.49 [1.35, 1.63] |
| | **Loss** | 4.96 [4.53 , 5.38 ] | 4.87 | 1.02 [0.89, 1.15] |
| **Reference** | **Gain** | 9.05 [8.37 9.7] | 7.8 | 1.64 [1.45 1.83] |
| | **Loss** | 6.80[6.11 7.5] | 7.84 | 1.23 [1.06 1.4 ] |
| **EEGLAB** | **Gain** | 6.1 [5.53 6.67] | 6.03 | 1.43 [1.24 1.62 ] |
| | **Loss** | 3.66 [3.12 4.2] | 5.7 | 0.91 [0.75 1.06] |
| **BrainStorm** | **Gain** | 8.43 [7.73 9.14] | 6.8 | 1.75 [1.52 1.98] |
| | **Loss** | 5.6 [4.98 6.34] | 6.57 | 1.21 [1.03 1.41] |
| **FieldTrip** | **Gain** | 7.32 [ 6.6 7.9] | 7.526961 | 1.37 [1.20 1.55] |
| | **Loss** | 4.67 [4.08 5.26] | 6.693929 | 0.98 [0.83 1.13] |

Table S2. The descriptive statistics and the effect size of the gain and loss conditional ERP, using the meak peak measure, for the compared softwares, with rejecting trials exceeding 200 microVolts min-max . The reported mean, standard deviation and cohen’s d values were computed across subjects.
